# Correlative SIP-FISH-Raman-SEM-NanoSIMS links identity, morphology, biochemistry, and physiology of environmental microbes

**DOI:** 10.1101/2022.03.29.486285

**Authors:** George A. Schaible, Anthony J. Kohtz, John Cliff, Roland Hatzenpichler

## Abstract

Microscopic and spectroscopic techniques are commonly applied to study microbial cells but are typically used on separate samples, resulting in population-level datasets that are integrated across different cells with little spatial resolution. To address this shortcoming, we developed a workflow that correlates several microscopic and spectroscopic techniques to generate an in-depth analysis of individual cells. By combining stable isotope probing (SIP), fluorescence *in situ* hybridization (FISH), scanning electron microscopy (SEM), confocal Raman microspectroscopy (Raman), and nano-scale secondary ion mass spectrometry (NanoSIMS), we illustrate how individual cells can be thoroughly interrogated to obtain information about their taxonomic identity, structure, physiology, and metabolic activity. Analysis of an artificial microbial community demonstrated that our correlative approach was able to resolve the activity of single cells using heavy water SIP in conjunction with Raman and/or NanoSIMS and establish their taxonomy and morphology using FISH and SEM. This workflow was then applied to a sample of yet uncultured multicellular magnetotactic bacteria (MMB). In addition to establishing their identity and activity, backscatter electron microscopy (BSE), NanoSIMS, and energy-dispersive X-ray spectroscopy (EDS) were employed to characterize the magnetosomes within the cells. By integrating these techniques, we demonstrate a cohesive approach to thoroughly study environmental microbes on a single cell level.

## Introduction

Efforts to understand the ecology of mixed environmental microbial populations are often hindered by our inability to collectively study the morphology, physiology, and taxonomy of distinct taxa on a meaningful scale. By correlating different single cell resolving spectroscopic and microscopic methods onto a single sample, more meaningful ecophysiological interpretations can be made. Cultivation-independent techniques such as microautoradiography and nano-scale secondary ion mass spectrometry (NanoSIMS), as well as next generation physiology approaches such as Raman microspectroscopy (Raman), have been used to study the *in situ* function and interactions of uncultured microbes (1). In conjunction with omic tools, these techniques provide insights into the chemical composition and metabolic activity of cells. However, because they are typically applied separately, meaningful correlations between individual cells and the relevant sets of measurements are often hard (or impossible) to draw. By applying multiple microscopic and spectroscopic analyses on a single sample, an approach referred to as correlative microscopy (2), the limitations of a single technique can be overcome, and the results combined into a single dataset that provides micro-to nano-scale information of a biological sample (3).

Correlative microscopy was developed as a means to overcome the limitations in resolution of conventional light microscopy by combining fluorescence microscopy (FM) with electron microscopy (EM), thus allowing for ultrastructural features within eukaryotic cells to be studied in greater detail (4, 5). The correlation of light and electron microscopy has since become a standard tool for the study of human cells and tissues (6-10), but is comparatively underdeveloped in microbial ecology. Recent approaches in microbial ecology have combined 16S rRNA targeted fluorescent *in situ* hybridization (FISH) with scanning electron microscopy (SEM) to study specific populations of magnetotactic bacteria that had been magnetically enriched from their respective habitats (11-13). Additionally, FISH has been used in combination with transmission electron microscopy (TEM) to study the ultrastructural differences between cells of uncultured methanotrophic consortia (14, 15) and with atomic force microscopy and Raman to study the morphology of cells belonging to the candidate phylum Acetothermia (16). An alternative approach, which has not yet been combined with FISH, is to resin-embed natural samples of appropriate water content (soil, sediment) and study the sample’s spatial structure and cell activity via a combination of FM, EM, energy-dispersive X-ray spectroscopy (EDS), and microtomography (17-19). Analytical techniques such as NanoSIMS have offered an additional dimension of information, providing nanometer resolution maps of the elemental and isotopic composition of cells (20). This technique has been used in conjunction with stable isotope probing (SIP) to map the distribution of labeled cellular products within eukaryotic cells and correlate it with FM and TEM (9). In addition to SIP-FISH-NanoSIMS, SIP-FISH-Raman has also been used to explore the activity of microbes (21, 22), though to our knowledge Raman has never been correlated with NanoSIMS and SEM. Despite these exciting developments, the use of correlative microscopy in microbial ecology has remained limited.

Here, we illustrate a workflow to correlate FISH, SEM, Raman, and, if desired, NanoSIMS to provide a comprehensive characterization of microorganisms at single cell resolution. This approach enables the study of the taxonomic identity (rRNA-targeted FISH), morphology (SEM), biochemistry (Raman), and metabolic activity (SIP-Raman and/or SIP-NanoSIMS) of cells within microbial communities or phenotypic heterogeneity within a clonal population of cells. This is, to our knowledge, also the first correlative application of both Raman microspectroscopy and NanoSIMS on the same cells.

We first tested our workflow on an artificially mixed community composed of the bacterium *Escherichia coli* and the archaeon *Methanosarcina acetivorans*. Next, we applied the correlative workflow to multicellular magnetotactic bacteria (MMB; *a*.*k*.*a*. multicellular magnetotactic prokaryotes, MMP), which are affiliated with the bacterial phylum Desulfobacterota (previously described as the class Deltaproteobacteria (23)) that are typically found in salty-brackish coastal habitats (24). MMB grow as single-species consortia composed of 15-60 cells that arrange themselves in a spherical or oblong shape around an acellular, central compartment (25-27). A hallmark of magnetotactic bacteria, such as MMB, is their ability to biomineralize ferrimagnetic crystals of magnetite (Fe_3_O_4_) and/or greigite (Fe_3_S_4_) and encapsulate them in lipid vesicles called magnetosomes (28-30). These organelles allow magnetotactic bacteria to orient themselves in Earth’s magnetic field, a phenomenon termed magnetotaxis, which can be exploited to magnetically enrich them from environmental samples. This is particularly important as MMB typically are rare community members and have proven recalcitrant to cultivation.

The MMB used in this study are found in the Little Sippewissett salt marsh (LSSM), a brackish marsh located in the state of Massachusetts (USA) (31, 32) that for decades has been used as a model system for microbial ecology. Although several studies on the microbiology of LSSM employed powerful visualization techniques, including FM, SEM, and NanoSIMS, none of these studies applied a correlative microscopic workflow and none applied it to MMB (19, 31-38). By applying our correlative workflow to the MMB found in LSSM, we were able to study the morphology and relative metabolic activity of three MMB populations that coexist in LSSM. Separately, we applied backscatter scanning electron microscopy (BSE-SEM) and EDS to image the magnetosomes of the MMB. Together with Raman and NanoSIMS these techniques were used to confirm the localization of iron (Fe) and sulfur (S) in the magnetosomes, suggesting that three of the five MMB populations found in LSSM use greigite as the ferrimagnetic mineral in their magnetosomes.

## Materials and Methods

### Preparation and stable isotope probing of an artificial community

A mock community was prepared by mixing cultures of *Escherichia coli* K12 (DSM498) and *Methanosarcina acetivorans* C2A (DSM2834) that had been grown in either the presence or absence of deuterated water (D_2_O). *E. coli* was grown aerobically with agitation (200 rpm) at 37 °C for 4 hours from an OD_600_ of 0.04 to an OD_600_ of 0.2 in either unamended M9 medium or medium that had been amended to a final concentration of 30% D_2_O using D_2_O (99.9%-D; Cambridge Isotope Laboratories). *M. acetivorans* cultures were grown anaerobically without agitation for 24 hours in DSMZ Medium 141c, either without D_2_O or amended to a final concentration of 30% D_2_O. At the end of the incubation, 1 mL of each culture was chemically fixed by adding paraformaldehyde (PFA; Electron Microscopy Science; EM grade) to a final concentration of 2% and incubating the cell suspension for 60 minutes at room temperature. Afterwards, cells were washed twice with 1x phosphate buffered saline (PBS; pH 7.4) by centrifugation at 16,000 *g* for 5 minutes, after which their supernatants were removed, and the cell pellets were resuspended in 1 mL of 1x PBS. A mock community was then made by mixing approximately equal number of cells from each suspension, resulting in a mixture of unlabeled and deuterium-labeled *E. coli* and *M. acetivorans* cells. Cells were stored at 4 °C in 1x PBS.

### Collection and stable isotope probing of environmental sample

Approximately 1 L of sediment slurry (7:3 sediment:water) was collected from a tidal pool in LSSM (41.5758762, -70.6393191) in Falmouth, MA (USA) during low tide on August 6^th^, 2021. In addition, 1 L of overlying marsh water was collected and filter sterilized using a 0.22 µm Millipore (Burlington, MA) Isopore PC filter for later preparation of SIP incubations. Within one day, samples were shipped on ice to Montana State University, Bozeman, MT (USA), where the sediment slurry was transferred to a 1 L glass beaker and stored in the dark for two days at ambient laboratory temperature (∼22ºC). MMB were enriched from the sediment by placing the South end of a magnetic stir bar against the exterior of the glass beaker just above the sediment layer, agitating the sediment by stirring, and then allowing the sediment to settle for 60 minutes. 200 µL of the water surrounding the accumulated magnetotactic bacteria was then removed with a pipette. The magnetic enrichment was repeated two additional times and all sampled liquids containing MMB were pooled into a single 1.5 mL tube containing 1 mL of 0.22 µM filtered marsh water. The sample was stored at room temperature for approximately 30 min before starting incubations with D_2_O. To incorporate D_2_O into the SIP incubations without changing the natural ionic composition of the sample, 200 mL of sterile-filtered marsh water were boiled until the volume was reduced to 100 mL, cooled to room temperature, and 100 mL of D_2_O was added for a final concentration of 50% D_2_O. The solution was purged with N_2_ for ten minutes to make it anoxic. To achieve a higher concentration of our target population in the sediment, MMB recovered by magnetic enrichment were inoculated into sealed serum vials containing 10 mL of 50% D_2_O salt marsh water. A live control sample was prepared by inoculating 100 µL of magnetically enriched MMBs into sealed serum vials with sterile-filtered marsh water without D_2_O. Headspace (∼15 mL) was replaced with N_2_ and samples were incubated for 24 hours at room temperature without agitation in the dark. After incubation, the serum vials were opened, and their contents emptied into 15 mL Corning Falcon tubes (Corning, NY). Magnetotactic bacteria were magnetically enriched from the 50% D_2_O salt marsh water and subsequently magnetically enriched two more times (10 minutes each) in 0.22 µm filtered marsh water before the samples were chemically fixed with 4% PFA for 60 minutes at room temperature. The samples were then centrifuged for 5 minutes at 16,000 g, after which the supernatant was removed, and the cell pellets resuspended in 50 µL 1×PBS and stored at 4 °C.

### Slide preparation

To successfully correlate different analytical methods, an appropriate surface substrate with minimal Raman background was needed. We used mirrored stainless steel due to its desired properties as a Raman substrate (39), its low cost, and it’s compatibility with both SEM and NanoSIMS. Circular coupons of mirror-finished 304 stainless steel (25 mm diameter, 0.6 mm thickness) were purchased from Stainless Supply (Monroe, NC). The coupons were cleaned by washing with a 1% solution of Tergazyme (Alconox, New York, NY) and rinsed with Milli-Q water, followed by sequential one-minute washes in acetone and 200 proof ethanol. Finally, the coupons were dried under compressed air and stored at room temperature. To maintain correct orientation of the samples, asymmetric boxes were etched into the mirrored surface of each coupon using a razor blade. 1 µL of each sample was spotted on the slide and air-dried at 46 °C for 1 minute, after which slides were washed by dipping them into ice-cold Milli-Q water to remove trace buffer components and air dried using compressed air. No further preparation was needed for Raman or SEM analysis.

### Confocal Raman microspectroscopy and spectral processing

Raman spectra of individual cells in the artificial community or MMB enrichments were acquired using a LabRAM HR Evolution Confocal Raman microscope (Horiba Jobin-Yvon) equipped with a 532 nm laser and 300 grooves/mm diffraction grating. Spectra of cells in the mock community and the MMB enrichments were acquired using a 100x dry objective (NA = 0.90) in the range of 200-3,200 cm^-1^, with 2-4 acquisitions of 10 seconds each, and a laser power of 4.5 mW. Spectra were processed using LabSpec version 6.5.1.24 (Horiba). The spectra were preprocessed with a Savitsky-Goly smoothing algorithm, baselined, and finally normalized to the maximum intensity within the 2,800-3,100 cm^-1^ region. To analyze the degree of deuterium substitution in C-H bonds (%C-D, *i*.*e*. (C-D_area_/(C-D_area_+C-H_area_)*100)), the bands assigned to C-D (2,040–2,300 cm^-1^) and C-H (2,800–3,100 cm^-1^) were calculated using the integration of the specified regions (21). Greigite (Fe_3_S_4_) was identified by its characteristic peak at 350 cm^-1^ (40). Neither magnetite (Fe_3_O_4_; 303, 535, and 665 cm^-1^ (40)), nor its laser-induced oxidative product, hematite (Fe_2_O_3_; 225, 245, 291, 411, and 671 cm^-1^ (40)), were observed.

### Scanning electron microscopy

To acquire SEM images of the mock community and MMB enrichments, a Zeiss (Jena, Germany) SUPRA 55VP field emission scanning electron microscope (FE-SEM) was operated at 1 kV under a 0.2-0.3 mPa vacuum with a working distance of 5 mm at the Imagining and Chemical Analysis Laboratory (ICAL) of Montana State University (Bozeman, MT). No conductivity coating was applied prior to SEM analysis due to the operation of the microscope at 1 keV.

### Backscatter electron microscopy and energy-dispersive X-ray spectroscopy

To image magnetosome minerals within MMB, backscatter electron microscopy (BSE) and energy-dispersive X-ray spectroscopy (EDS) were used. The ICAL Zeiss SUPRA 55VP FE-SEM was operated at 10 kV with a working distance of 7.5 mm and an aperture of 30 µm. For EDS, an accelerating voltage of 10 kV was used under high current and the aperture set to 60 µm. EDS data was collected using an Oxford Instruments (Abingdon, UK) AZtec detector with a pixel resolution of 2,048 × 2,048 and a pixel dwell time of 100 µs. Qualitative elemental abundances of C, N, O, Si, P, S, Ca, Cr, Mn, Fe, and Ni were acquired, and data were processed using the AztecLive software (Oxford Instruments, Abingdon, UK). Elemental maps of C and S were used to identify the location and orientation of the magnetosomes within the MMB. Iron was not used to identify the magnetosomes due to the overwhelming iron signal from the stainless steel coupon.

### Fluorescence in situ hybridization

Double-labeled oligonucleotide probes for FISH (DOPE-FISH) (41) were purchased from Integrated DNA Technologies (Coralville, IA) to visualize different taxa by FM. Cells were dehydrated using an increasing ethanol series (1 min in each 50, 80, and 96% ethanol) and 16S rRNA-targeted FISH was carried out directly on the stainless steel coupon following established protocols (42). Samples were hybridized using either FAM-, Cy3-, or Cy5-labeled DOPE-FISH probes for 2 h (artificial community) or 3 h (MMB) in a humid chamber at 46 °C at a final probe concentration of 2.5 ng µl^-1^. Probe mix EUB338 I-III and ARC915, targeting most bacteria and archaea respectively, were used at 35% formamide (43, 44).

Three newly designed DOPE-FISH probes targeting the 1,032-1,048 nt region of the 16S rRNA (*E. coli* equivalent) were applied to the MMB enrichments using previously published 16S RNA gene clone sequences (32, 45) as well as currently unpublished metagenomic 16S rRNA gene sequences (Schaible *et al*., unpublished). These probes target three populations of MMB in LSSM, Group 1 (G1MMB1032; FAM-5′-CCTGTCATCGGGCTCCCC-3′-FAM; Tm = 60.8 °C), Group 3 (G3MMB1032; Cy3-5′-CCTGTCTTTGGGCTCCCC-3′-Cy3; Tm = 58.4 °C), and Group 4 (G4MMB1032; Cy5-5′-CCTGTCTTCAGGCTCCCC-3′-Cy5; Tm = 57.8 °C) and were used at 50% formamide. Hybridizations were performed using an equimolar mixture of the three probes and two competitor probes targeting the same region of the 16S rRNA to increase the stringency of the hybridization, cG2MMB1032 (5′-CCTGTCATCGGGTTCCCC-3′; Tm = 58.1 °C) and cG5MMB1032 (5′-CTTGTCTTCAGGCTCCTC-3’; Tm = 52.3 °C). Because MMB groups 2 and 5 were of low abundance at the time of sampling (<1% of all MMB), these populations were not targeted in the workflow. Hybridizations with probe NonEUB338-I (FAM-5’-ACTCCTACGGGAGGCAGC-3’-FAM) were used as negative controls (46).

### Nano-scale secondary ion mass spectrometry

Prior to NanoSIMS data acquisition, sample regions of interest (ROIs, ∼150 × 150 µm) were marked on the stainless steel surface using a Leica LMD6 Laser Microdissection System (Wetzlar, Germany). To spatially map the elemental composition and the relative isotopic abundances of hydrogen of our mock community and MMB enrichments, ion images were acquired using the NanoSIMS 50L (Cameca) at the Environmental Molecular Sciences Laboratory at the Pacific Northwest National Laboratory. All NanoSIMS images were acquired using a 16 keV Cs^+^ primary ion beam at 512 × 512-pixel resolution with a dwell time of 13.5 ms px^-1^. Analysis areas were pre-sputtered with ≥10^16^ ions cm^-2^ prior to analysis. D^-^ and H^-^ secondary ions were accelerated to 8 keV and counted simultaneously using electron multipliers (EMs). The vacuum gauge pressure in the analytical chamber during all analyses was consistently less than 3×10^−10^ mbar. Other analytical conditions included a 200 µm D1 aperture, 30 µm entrance slit, 350 µm aperture slit, and 100 µm exit slits. Secondary tuning was adjusted, and D^-^ and H^-^ peaks were monitored between analyses for drift. The OpenMIMS plugin for ImageJ was used to access and correct images pixel by pixel for dead time (44ns) and QSA (β=0.5). Data from ROI were exported to a custom spreadsheet for data reduction. Semi-quantitative D/H analyses were calibrated against an in-house yeast reference material of unknown, but natural abundance δD during the same analytical session using similar conditions to those used to analyze the bacterial culture samples. The yeast reference material had been stored in the NanoSIMS under high vacuum for several months prior to the analyses reported here. Hydrogen isotope analyses were acquired using a 4-pA primary beam yielding a primary beam size of about 150 nm. Detectors collecting H^-^ and D^-^ ions were situated near the center of the magnet radius to improve simultaneous secondary centering characteristics. Hydrogen isotopes are reported as δD relative to the yeast standard and (uncalibrated) apparent atom %. Propagation of uncertainty includes counting statistics and external precision of D/H ratios of 16 individual yeast cells. To explore colocation of Fe and S atoms in the MMBs, some images were acquired by counting ^32^S^-^ and ^32^S^56^Fe^-^ ions simultaneously using a 2-pA primary beam.

## Results and Discussion

### General considerations for correlative microscopy

Correlative imaging enables identification and characterization of diverse microbial taxa on a single cell level, allowing for a complimentary suite of correlated data to be collected. Prior to sample analyses, the abundance, morphology, and fixation of the target cells need to be considered to ensure that the desired population of interest is studied with its morphology and isotope ratios intact. The ratio of fixatives may need to be optimized on a sample-specific basis and should be tested prior to the correlative workflow. For example, McGlynn *et al*. 2018 applied several fixation protocols optimized for specific visualization goals, using a mix of PFA and glutaraldehyde at varying concentrations (14). Here, cell fixation was performed using 2% PFA (*E. coli* and *M. acetivorans*) and 4% PFA (MMB) because these conditions were found to maintain cellular structure without autofluorescence from the fixative, which can be an issue with glutaraldehyde (14).

Depending on the relative abundance and morphology of the target population, our correlative workflow can be employed either FISH-first or SEM-first (Fig. 1). Microbes with common morphologies, such as coccoid, rod, or filament shapes, or those present at low relative abundances, will likely need to be identified using FISH before other analyses are employed to ensure the correct phylotype or morphotype is studied (12). Cells with distinct morphologies, or cells that are present at high relative abundance, can be imaged by SEM first, followed by Raman and FISH. Alternatively, the target organism can be physically enriched prior to downstream analyses. For example, here we magnetically enriched a lowly abundant population of multicellular magnetotactic bacteria (0.003%-0.15% relative abundance in LSSM according to 16S rRNA gene amplicon data ((19) and Schaible and Hatzenpichler unpublished). An alternative is to separate FISH-labeled cells via fluorescence-activated cell sorting, as was previously shown (47, 48). An SEM-first workflow is advantageous when cells or cell aggregates, such as the MMB studied here, are susceptible to changes in morphology triggered by the repeated dehydration-rehydration cycles used in FISH protocols (Fig. S1).

**Figure 1.**
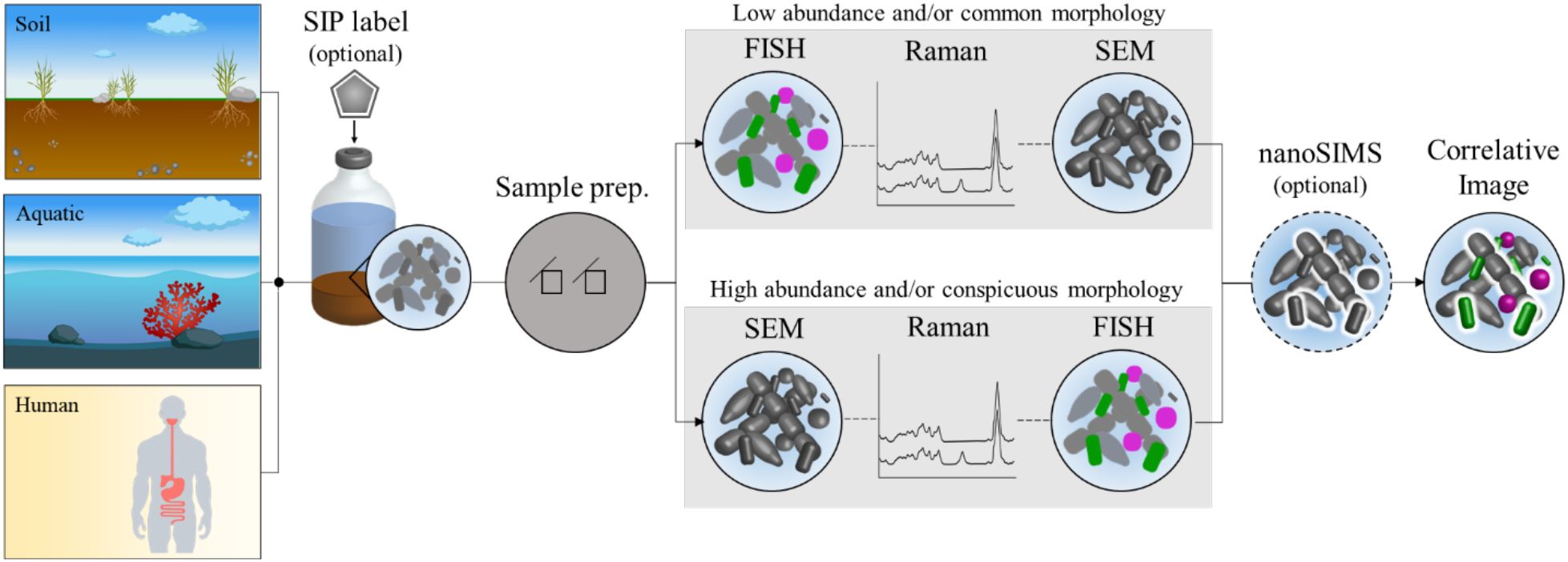
Correlative microscopy workflow. Environmental samples can be taken and incubated in the presence of an isotope-containing substrate to label active community members or study substrate assimilation. Next, the biomass is chemically fixed and placed on a stainless steel coupon. SEM is used to study cell morphology. Raman is used to determine the biochemical makeup as well as substrate assimilation of individual cells. rRNA-targeted FISH reveals the taxonomic identity of cells. As a final step, NanoSIMS can be used to study the elemental and isotopic composition of the sample at higher spatial resolution and sensitivity than possible by Raman.

Sample treatment prior to NanoSIMS has been shown to decrease the isotope enrichment, resulting in underestimates of activity (49). For this reason, it is important to limit treatments to the sample during preparation and reduce unnecessary exposure to or inclusion of exogenous material during these treatments that could lead to dilution of the isotopic label (49). Furthermore, the use of mono-FISH instead of catalyzed reporter deposition fluorescence *in situ* hybridization (CARD-FISH) has been recommended to limit dilution of the isotope signal due to introduction of the horseradish peroxidase conjugated probes and/or excessive tyramide deposition (49, 50). While the use of mono-FISH has been recommended, poor fluorescent signal due to, for example, low ribosomal content in slow-growing cells, may necessitate the use of CARD-FISH (51). In this study, we used DOPE-FISH (two fluorophores) instead of mono-labeled FISH probes to increase fluorescence yield (41).

### FISH-first correlative workflow using an artificial microbial community

To benchmark our FISH-first correlative workflow, we prepared an artificial community by mixing *E. coli* and *M. acetivorans*, both representing common morphotypes (rods and cocci, respectively). Cultures were grown in the presence or absence of deuterated water (D_2_O) as a general marker of anabolic activity (21). PFA-fixed biomass was immobilized on a stainless steel coupon that had been etched with a razor blade to allow tracking of the same area throughout all analyses (Fig. S2). We then applied 16S rRNA-targeted DOPE-FISH (41) to identify bacterial and archaeal cells and choose specific regions of interest (ROIs) for downstream analysis (Fig. 2a). Following DOPE-FISH, Raman was used to study the chemical composition of cells in the same ROIs (Fig. 2b, Table S1). Prior to acquisition of Raman spectra, cells were photobleached for 1-2 minutes to remove background from the FISH dyes (52). Alternatively, dyes such as FAM/FITC or Cy5, that are not excited by the 532 nm Raman laser, could be used (22).

**Figure 2.**
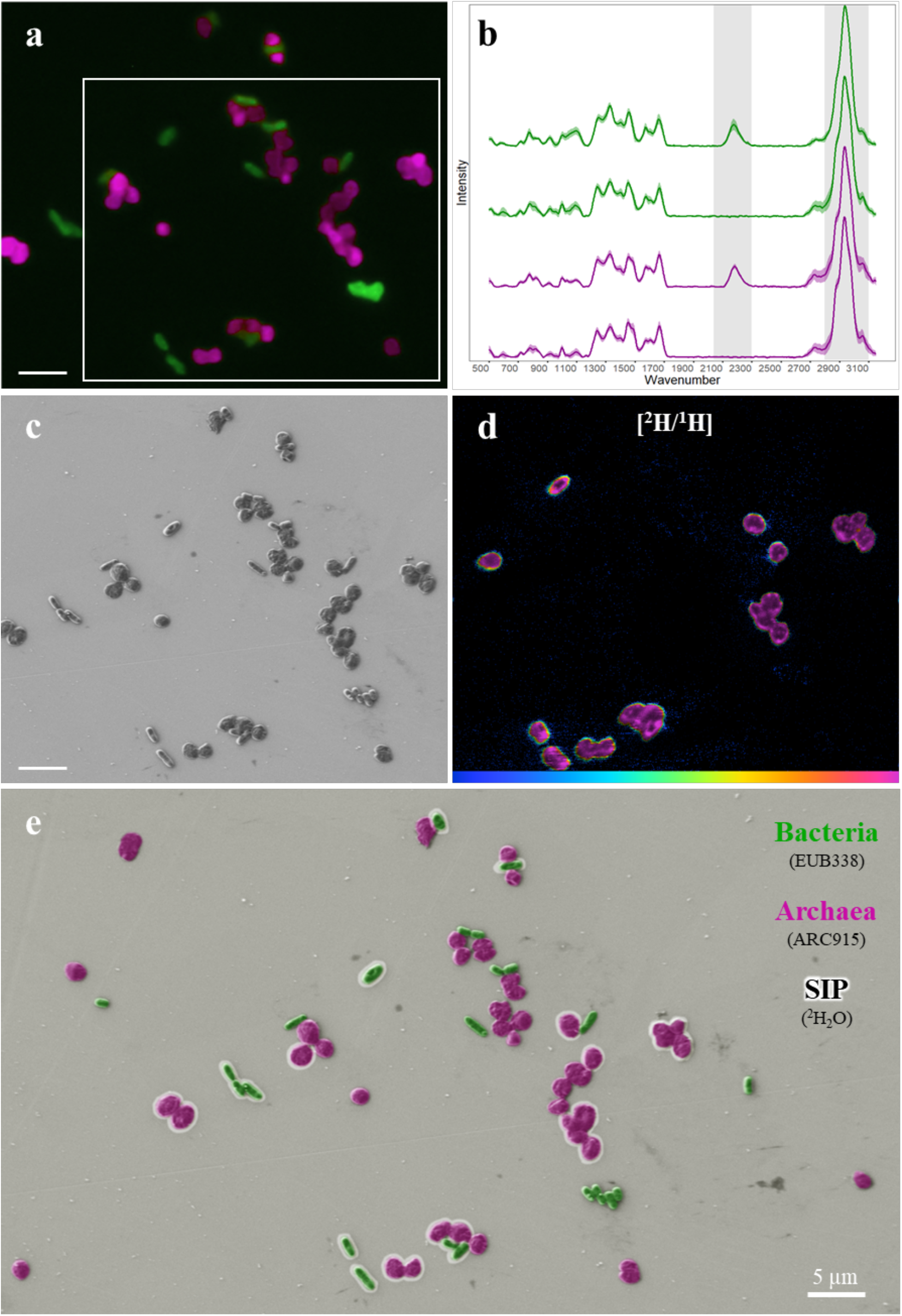
Correlative imaging of an artificial community. (a) FISH targeted *E. coli* (EUB388mix, green) and *M. acetivorans* (ARC915, magenta). (b) Single cell Raman identifies cells that had taken up deuterium from D_2_O (detected by the characteristic peak shifts of C-H at 2,800-3,100 cm^-1^ to C-D at 2,040-2,300 cm^-1^, highlighted in gray). Each spectrum shown is an average of the cells shown in panel a with the shaded regions showing the standard deviation for each data set. The green spectra correspond to D_2_O positive (top; n = 4) and negative (bottom; n = 10) *E. coli* cells. The magenta spectra correspond to D_2_O positive (top; n = 14) and negative (bottom; n = 16) *M. acetivorans* cells. (c) Cell morphology revealed by SEM. (d) NanoSIMS hue-saturation-intensity image of ^2^H/^1^H ratio confirms deuterium uptake at higher spatial resolution and sensitivity than Raman. The area analyzed using NanoSIMS is outlined in panel a. The scale is 0 – 2.0 atom %. (e) Composite false color image presenting all data. Cells outlined by a white halo were deuterium-labeled as determined by both Raman and NanoSIMS as both methods yielded consistent results. Cells without a halo were considered unlabeled. All scale bars equal 5 µm.

The incorporation of deuterium into biomass was calculated using the integration of the C-D (2,040-2300 cm^-1^) and C-H (2,800-3,100 cm^-1^) regions of the spectra (Fig. 2b; (21)). This showed that labeled *E. coli* and *M. acetivorans* had an average of 11.0% (± 2.5) and 9.3% (± 1.4) C-D (*i*.*e*., (CD/(CD+CH))*100), respectively, and that both unlabeled *E. coli* and *M. acetivorans* had an average of 0.5% (± 0.3 and 0.4, respectively) C-D (Fig. S3, Table S1). Next, field emission SEM (FE-SEM) was performed to reveal cell morphology (Fig. 2c). After FE-SEM images were taken, specific ROIs were traced using a laser dissection microscope to assist in finding imaged areas during NanoSIMS analyses (Fig. S2c). This was necessary because SIMS instruments, such as the Cameca NanoSIMS 50L employed in this study, only offer a low magnification objective that makes finding the targets of choice difficult. Finally, the elemental and isotopic composition of the sample was imaged using NanoSIMS, which provided a map of the metabolic heterogeneity of the mixed population at single cell resolution (Fig. 2d, Table S1). Analysis of the NanoSIMS data revealed that the labeled *E. coli* and *M. acetivorans* had an average deuterium content of 5.2% (± 1.2) and 5.3% (± 1.1), respectively, and that both unlabeled *E. coli* and *M. acetivorans* showed no deuterium incorporation (Fig. S3, Table S1).

While Raman can provide information on cellular isotope uptake, its sensitivity is 30-100x lower than NanoSIMS, depending on the isotope (1, 53). Furthermore, the expression of %C-D (relative change from C-H to C-D stretch in lipids and proteins) cannot be interpreted as %D content of the entire biomass. In contrast, NanoSIMS is a destructive but fully quantitative technique capable of measuring the elemental and isotopic composition of a cell at atomic resolution. However, NanoSIMS does not provide reliable information about the molecular composition of a sample, which is, in turn, achieved by Raman (1, 52). These fundamentally different working mechanisms are a major reason for the discrepancy between the H/D ratios determined via Raman (11 %C-D for *E. coli*) vs. NanoSIMS (5.2 atom % D). Depending on the exact research question, extent of metabolic activity, and available instruments, researchers need to decide whether the use of NanoSIMS and/or Raman is warranted (54). Eventually, the information gained by these different analyses (*i*.*e*., taxonomy, morphology, and metabolic activity) can be projected into a single image that correlates each analysis (Fig. 2E). Both Raman and NanoSIMS data showed significantly different isotope enrichment between positively and negatively labeled cells (p-value = 3.87×10^−16^ and 4.23×10^−12^, respectively, t-test; Fig. S3, Table S1). Our analysis of the mock community provided a successful proof of concept.

The application of Raman and NanoSIMS on the same cells revealed a discrepancy between the two methods in detecting deuterium in the mock community (Fig. S4). The only other study, to our knowledge, that applied both Raman and NanoSIMS reported a similar trend of deuterium incorporation with respect to the deuterated water percent in the growth medium (21); however, their analysis was not performed on the same cells (*i*.*e*., uncorrelated data), which limits comparability between their and our datasets. Nevertheless, our newly developed workflow has highlighted a need for further study on the quantitative measure of isotope incorporation via Raman and NanoSIMS and to what extent results obtained via these two methods can be compared.

### SEM-first correlative workflow using environmental MMB

We established an SEM-first correlative workflow to interrogate MMB enriched from LSSM. Correlative fluorescence microscopy (FM) and SEM have previously been performed on both single celled magnetotactic bacteria (11, 12, 15) as well as MMB from the Mediterranean Sea (13), but to our knowledge the combined techniques have not been applied to MMB from LSSM. In addition, we show the first application of Raman and NanoSIMS to MMB. Based on 16S rRNA gene studies, the sample site at LSSM contains several distinct populations of MMB that have not been studied on a population-specific morphological and physiological level (19, 31, 32). To identify whether these populations differ in morphology and relative metabolic activity, we applied our new correlative microscopy workflow to MMB. MMB magnetically enriched from LSSM tidal pool sediment were incubated with 50% D_2_O to determine if MMB populations differ in their anabolic activity.

Because MMB exhibit a unique multicellular morphology, they can be easily distinguished from other morphotypes by light microscopy or SEM without the need for prior identification via FISH. This was of particular importance as MMB from LSSM suffer detrimental effects to their cellular morphology when subjected to FISH (Fig. S1). For this reason, FE-SEM was performed first on MMB, and specific ROIs identified (Fig. 3a). Raman was subsequently used to map the biochemical makeup of each MMB identified by FE-SEM. Raman spectra of MMB contained characteristic peaks for greigite (Fe_3_S_4_; 350 cm^-1^; (40)), indicating it is the mineral used in their magnetosomes (Fig. 4d). This is consistent with the absence of characteristic peaks for magnetite and its laser-induced oxidative product, hematite, in the Raman spectra (225, 245, 291, 411, and 671 cm^-1^ (40)). As done with the mock community, we attempted to identify the C-D peak in the silent region of the spectra (2,040-2,300 cm^-1^). Unexpectedly, we were unable to observe a C-D peak in any of the MMB spectra (Fig. 3b, Fig. S3, Table S1). At the time, we attributed this to a low labeling rate that was below the detection limit of Raman; this hypothesis was later confirmed by NanoSIMS (discussed below). Next, 16S rRNA targeted DOPE-FISH was used to identify three distinct MMB populations in LSSM (Fig. 3b) (32). As was done with the artificial community, ROIs were traced using a laser dissection microscope and the elemental and isotopic composition of the sample imaged using NanoSIMS (Fig 3d, Table S2).

**Figure 3.**
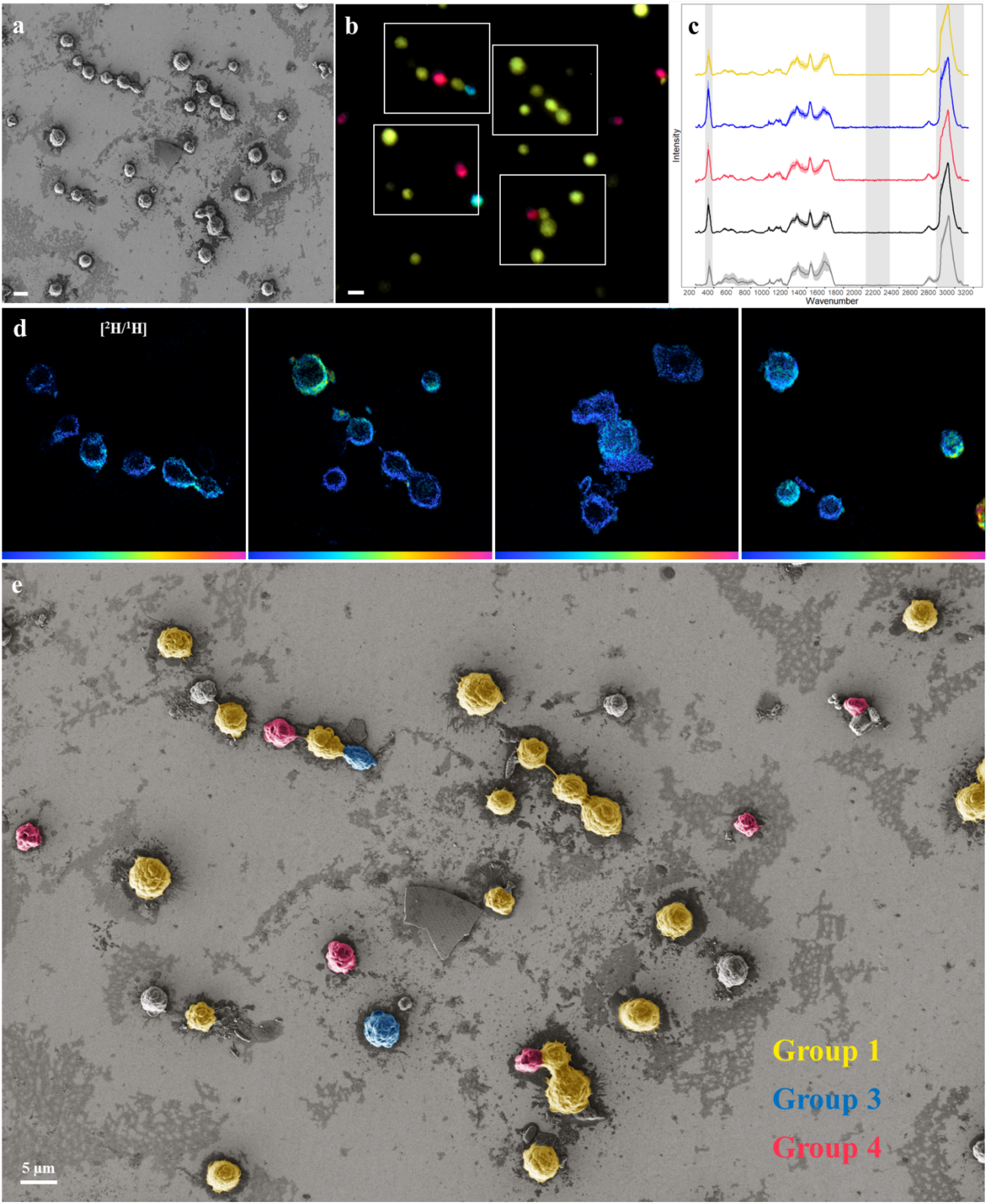
Correlative microscopy analysis of MMB. (a) SEM image of magnetically enriched MMB. (b) FM identifies three MMB subpopulations using FISH probes specific for Group 1 (yellow), Group 3 (teal), and Group 4 (red). Unlabeled MMB likely belong to two populations not targeted with FISH. (c) Raman spectra of MMB separated by the respective FISH label and control. The characteristic regions of C-H at 2,800-3,100 cm^-1^, C-D at 2,040-2,300 cm^-1^, and greigite at 350 cm^-1^ are highlighted in gray. Each spectrum shown is an average with the shaded regions showing the standard deviation for each data set. Raman spectra were collected from several ROIs besides the one shown in this figure. The yellow spectra show Group 1 (n = 38), the blue spectra show Group 3 (n = 2), and the magenta spectra show Group 4 (n = 3) MMB. The dark gray spectra are those of MMB that were not labeled by FISH (n = 10) and the light gray spectra show MMB from the negative control (*i*.*e*., no D_2_O addition; n = 27). (d) NanoSIMS hue-saturation-intensity images showing the ^2^H/^1^H ratios present in the ROIs outline in white boxes in b (scale is 0 to 0.55 atom %). (e) Composite false color image presenting data. MMB that are not false-colored were not labeled by one of the three FISH-probes. All MMBs were labelled with deuterium. All scale bars equal 5 µm.

**Figure 4.**
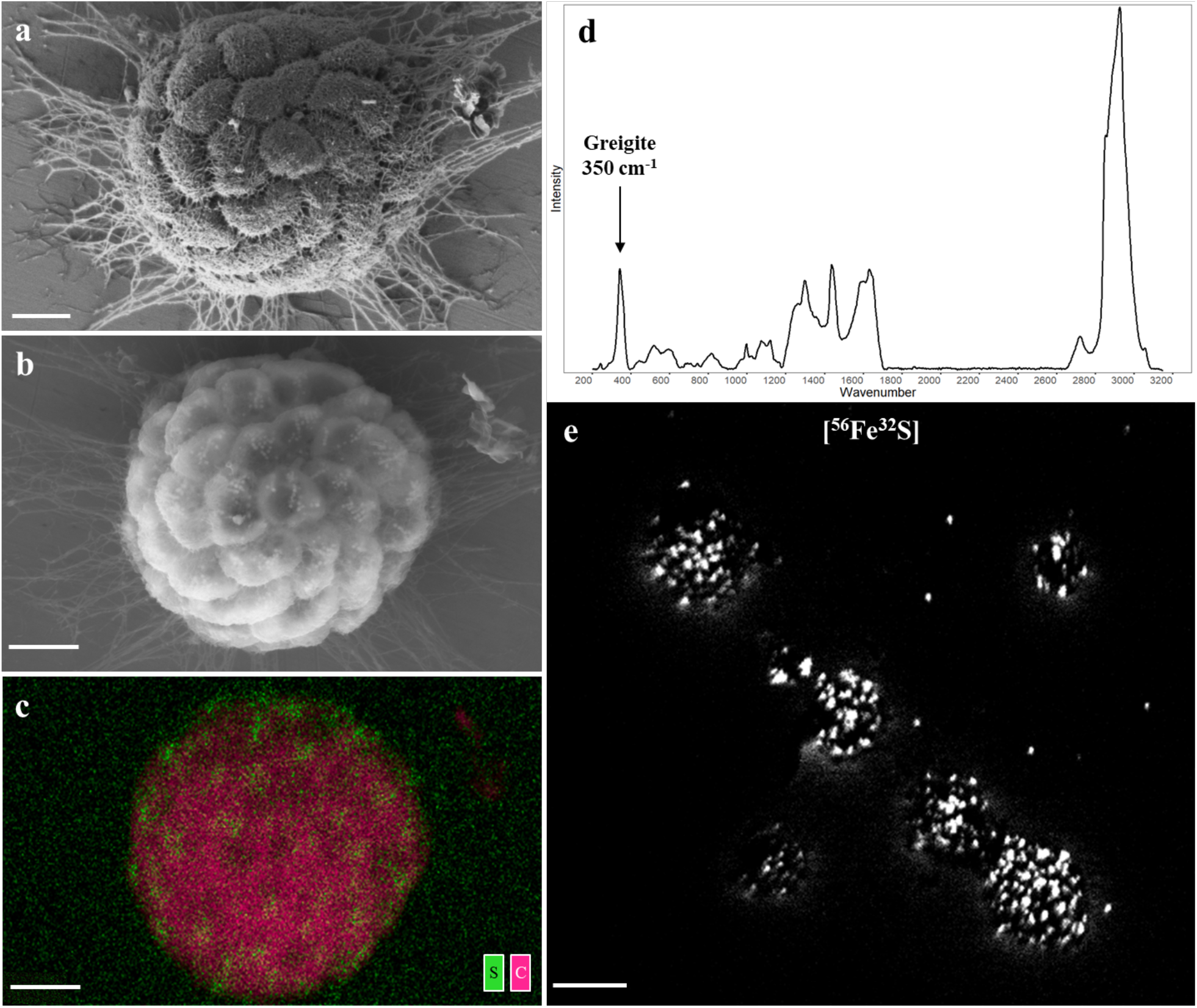
Imaging magnetosomes within MMB. (a) Low voltage SEM image of a MMB consortium. (b) BSE-SEM image of the same MMB reveals the magnetosomes in each cell seen as small white chains. (c) EDS image showing the localization of sulfur accumulations (green), presumably contained within magnetosome chains composed of greigite (Fe_3_S_4_), against cellular carbon (magenta). Fe and O are not shown because the stainless steel surface contains large amounts of Fe and O. (d) Raman spectra of a single MMB showing the characteristic peak for greigite (Fe_3_S_4_) at 350 cm^-1^ that was observed in all MMB analyzed (see Fig. 3c) (40). (e) NanoSIMS image showing the mass corresponding to ^56^Fe^32^S^-^ of the same MMB shown in the same ROI shown in Fig. 3d., indicating the localization of Fe and S within MMB. Panels a-c scale bars equal to 1 µm, panel e scale bar is equal to 5 µm.

While deuterium incorporation from D_2_O could not be detected via Raman, NanoSIMS showed that all MMB analyzed (n = 23) exhibited low levels of deuterium incorporation (0.11 - 0.78%; Table S1), significantly higher than the negative control (0.015 - 0.022%; p-value = 3.37×10^−19^; Fig. S3). While such values are indicative for deuterium incorporation, they are well below the detection limit of Raman (0.2% for D; (1)). Deuterium uptake from D_2_O has been shown to be affected by the organic carbon source used by heterotrophic cells for growth (21, 55). This makes it possible that the MMB studied here used organic carbon sources that contributed to the dilution of deuterium in cells. Nevertheless, such a low level of deuterium-incorporation from heavy water is surprising given that all MMB must have been metabolically active at the beginning and end of the SIP-incubation; otherwise, they would not have been able to swim to the magnetic stir bar used to collect MMB (which is an active process, they are not merely passively “pulled” to the magnet). As discussed above, it is possible that the fixation and FISH protocols used here resulted in an additional decrease in the observed isotope enrichment (49). Furthermore, it is possible that the level of deuterium in MMB was decreased during magnetic enrichments. If magnetotaxis is energetically expensive, enrichments of MMB in water lacking deuterium may have increased the D/H turnover and led to an overall decrease of deuterium in MMB cells. Future experiments will have to reconcile this conundrum.

The three MMB populations exhibited similar morphology when studied via FE-SEM, indicating that different MMB species found in LSSM cannot be distinguished by morphology alone (Fig. 3e), highlighting the importance and power of our correlative microscopy approach. Individual populations of MMB were identified using FISH, which revealed that Group 1 is the most abundant MMB population in our sample (66%, n = 64). Group 4 was the second most abundant (7%, n = 7), and Group 3 was the third most abundant (4%, n = 4). Unlabeled MMB accounted for 23% (n = 22) of all MMB and likely belonged to Groups 2 and 5 that were not targeted in this study. Our findings on the relative abundance of different MMB populations are consistent with the results reported by Simmons *et al*., who studied an unspecified tidal pool in LSSM (32).

### Analysis of MMB magnetosomes using BSE-SEM and EDS

Magnetotactic bacteria are relevant to the biogeochemical cycling of iron and sulfur, and the process of controlled biomineralization via magnetosome formation (30, 56). Historically, BSE and EDS have been used to study the location, shape, and elemental composition of the biogenic minerals within MMB (57-60). We applied BSE-SEM at an accelerating voltage of 10 kV, allowing for the electron-dense minerals within the magnetosomes of MMB to be imaged (Fig. 4b). EDS was then used at the same voltage to map elements across the cell, revealing confined areas of sulfur within individual cells (Fig. 4c). This result is consistent with the identification of greigite (Fe_3_S_4_) in the Raman spectra of MMB (Fig. 4d) as well as the co-localization of ^56^Fe and ^32^S, as demonstrated by the detection of ^56^Fe^32^S^-^ ions, observed via NanoSIMS (Fig. 4e) (40, 61). These findings align with observations that greigite is used as the ferrimagnetic mineral by MMB in other locations (57, 58, 62, 63).

Both BSE-SEM and EDS were applied outside of the correlative workflow due to the negative effect of high voltage electron beams on cells. MMB that had been analyzed using BSE-SEM and EDS could not be labeled using FISH, and the respective Raman spectra were consistent with carbonized biomass, likely due to the effect of using a high voltage electron beam (not shown). Previous correlative studies have used low dose electron imaging, such as FE-SEM, to avoid irreversible damage to the cell caused by high electron voltages (11).

## Conclusions

Understanding the metabolic potential and ecological distribution of microbial communities has been greatly aided by various omic techniques, but information on the morphology, physiology, and taxonomy of distinct taxa is lost during these analyses. Here, we provide a single cell level workflow to link the taxonomic identity and morphology of microbes using rRNA-targeted FISH and SEM with their cellular chemistry and metabolic activity using SIP-Raman and/or SIP-NanoSIMS. By applying this workflow, naturally low abundance populations (*e*.*g*., MMB) can be taxonomically identified and interrogated in greater detail than current practices allow. Furthermore, having correlated data is desirable for population studies to determine if physiological heterogeneity exists within clonal groups of cells.

In this study, we used D_2_O as a general marker of anabolic activity, but substrates labeled with ^13^C, ^15^N, ^18^O, or ^34^S could be incorporated in the workflow. Additionally, future applications and developments of this workflow might include the use of substrate analogue probing (1), super resolution microscopy (64, 65), or atomic force microscopy (16). Because substrate analogue probing labels specific biopolymers, such as proteins (66), nucleic acids (67), or lipid membranes (68), its use with traditional FM or super resolution microscopy can inform on overall biosynthetic activity or specific enzymatic function. Furthermore, correlative FISH-TEM approaches adapted from McGlynn *et al*. 2018 (14) and protocols to thin-section MMB developed by the group of Lins (27, 69) could provide valuable complementary information of the physiology and ultrastructure of the diverse MMB populations in LSSM.

## Author contributions

GAS, AJK, and RH designed the study. GAS and AJK optimized and tested the workflow. AJK performed the incubations and collected Raman spectra for the mock community. GAS performed the incubations and collected Raman spectra for the MMB. GAS performed FISH, SEM, BSE, and EDS. JC performed NanoSIMS analyses. GAS, AJK, and JC analyzed the data. GAS and RH wrote the paper. All authors reviewed and edited the paper.

## Acknowledgements

This study was funded through NASA FINESST award 80NSSC20K1365 (to G.S. and R.H.), NASA Exobiology award NNX17AK85G (to R.H.) and Gordon and Betty Moore Foundation award 5999 (to R.H.). Montana State University’s Confocal Raman microscope was acquired with support by the National Science Foundation (DBI-1726561) and the M.J. Murdock Charitable Trust (SR-2017331). This work was performed in part at the Montana Nanotechnology Facility, an NNCI member supported by NSF grant ECCS-2025391. A portion of this research was performed under the Facilities Integrating Collaborations for User Science (FICUS) program (proposal: 10.46936/fics.proj.2017.49972/6000002) and used resources at the Environmental Molecular Sciences Laboratory (https://ror.org/04rc0xn13), which is a DOE Office of Science User Facilities operated under Contract No. DE-AC05-76RL01830 (EMSL). We thank Jeff Marlow, Virginia Edgcomb, and Emil Ruff for help with collection of LSSM sediment samples, and Ashley Cohen and Zackary J. Jay for helpful comments on the manuscript.

## Description of Supplementary Files

### File Name

Supplemental Table 1

### Description

Table of the artificial mock community Raman and NanoSIMS data highlighting the output for single cells shown in Fig. 2. Because of the limited field of view possible by NanoSIMS, not every cell analyzed using Raman was also analyzed using NanoSIMS. Both deuterium (D)- labeled and unlabeled cells are shown in the table. EC, *Escherichia coli*. MA, *Methanosarcina acetivorans*.

### File Name

Supplemental Table 2

### Description

Table showing all Raman and NanoSIMS data used for MMB analyses. Because of the limited field of view possible by NanoSIMS in a single image, not every cell analyzed using Raman was also analyzed using NanoSIMS. G1, G3, G4 refer to FISH probes specific for MMB groups 1, 3, and 4, respectively.

**Supplemental Figure 1.**
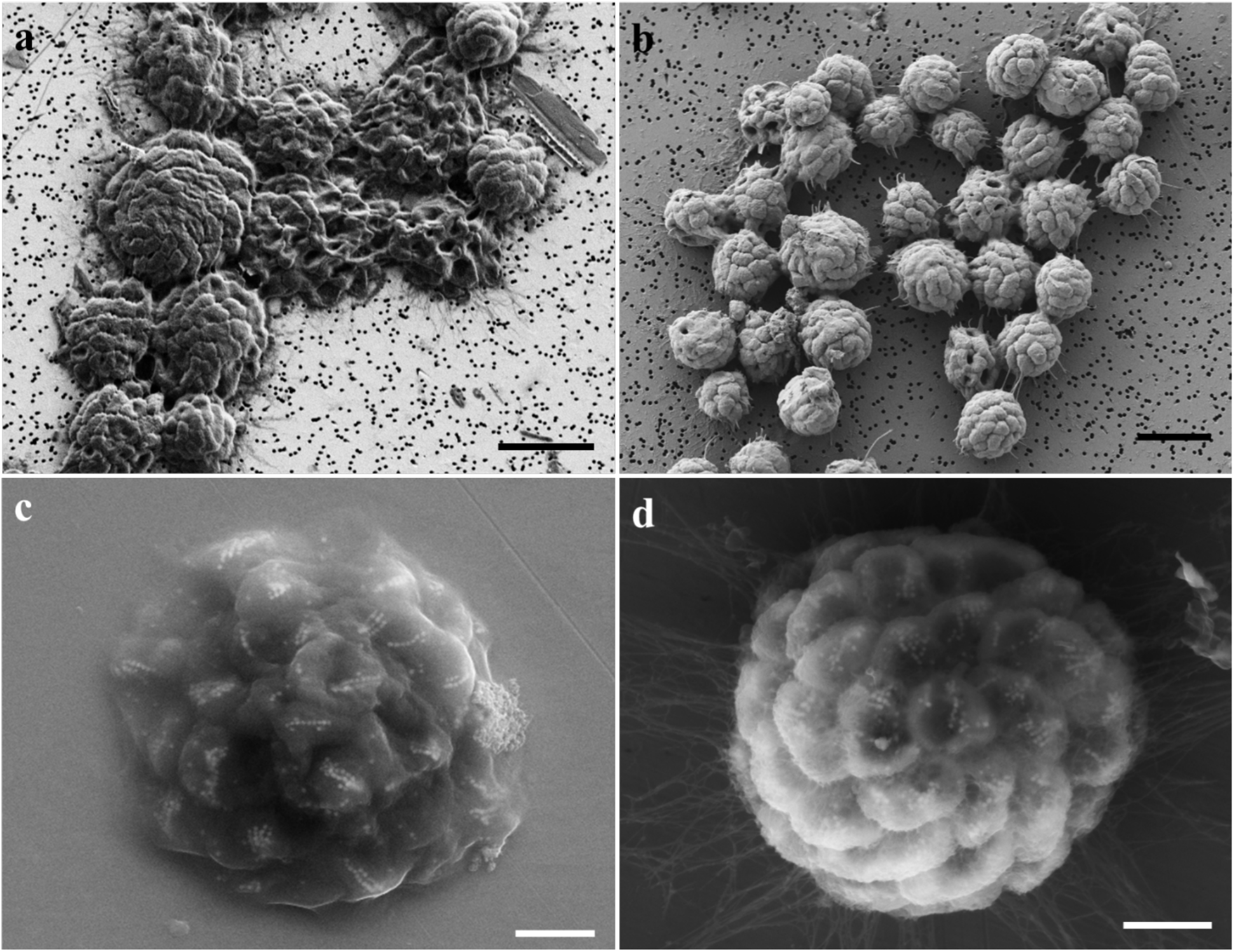
SEM images showing the loss of structural integrity of MMB when FISH is performed prior to SEM. (a-b) MMB deposited on a 0.22 µm filter and imaged using SEM (a) post and (b) prior to FISH. (c-d) MMB on a stainless steel coupon imaged using back scatter electrons with the secondary electron detector (c) post and (d) prior to FISH. Scale bars in a and b equal to 5 µm, c and d equal to 1 µm.

**Supplemental Figure 2.**
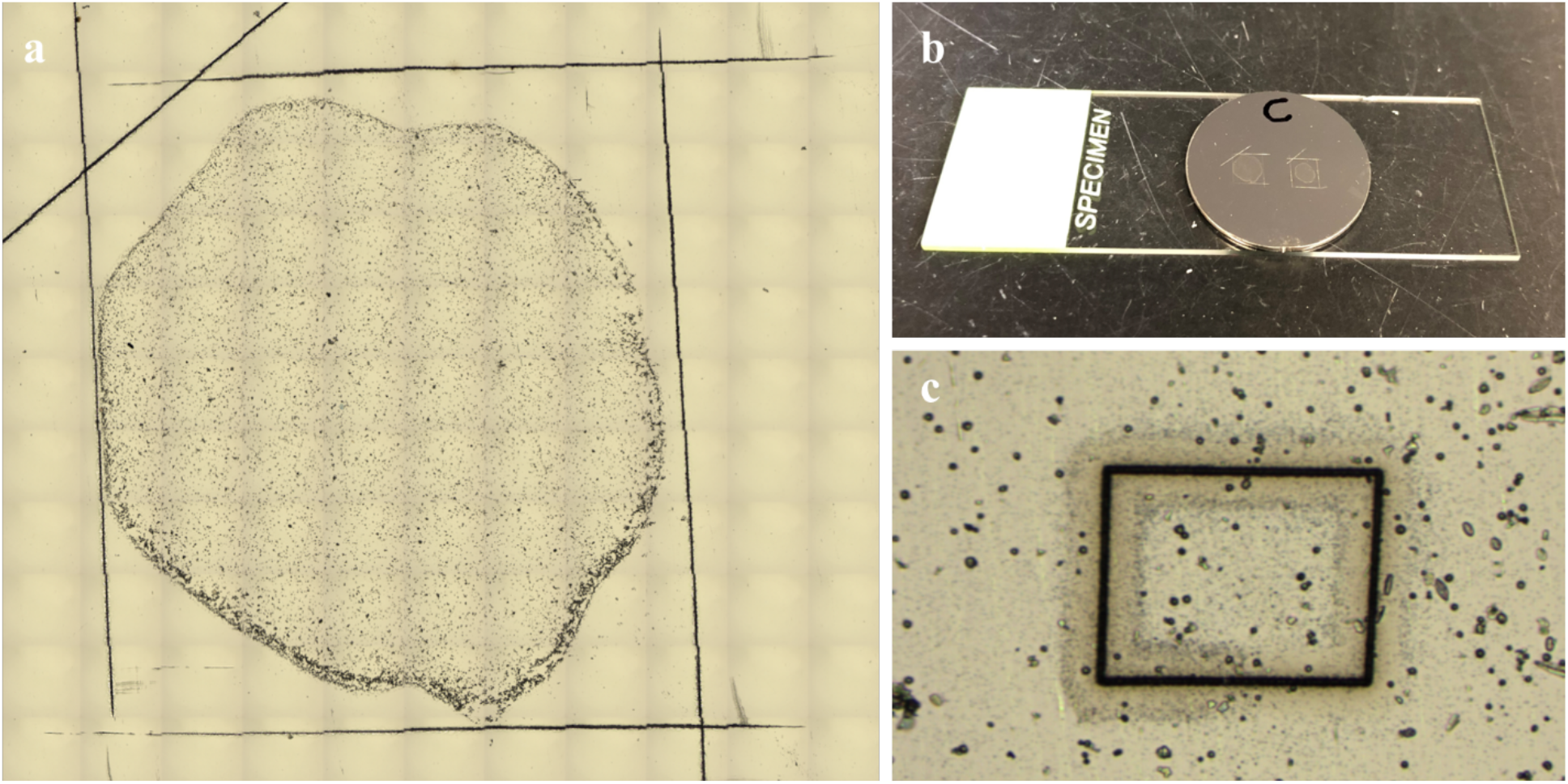
Slide design and sample orientation. (a) A mosaic image showing the sample dried within an asymmetric square etched into the stainless steel coupon with a razor blade. (b) The stainless steel coupon attached to a standard 60×20 mm microscope slide. (c) An ROI that had been traced using a laser dissection microscope to assist in locating the ROI during nanoSIMS analysis.

**Supplemental Figure 3.**
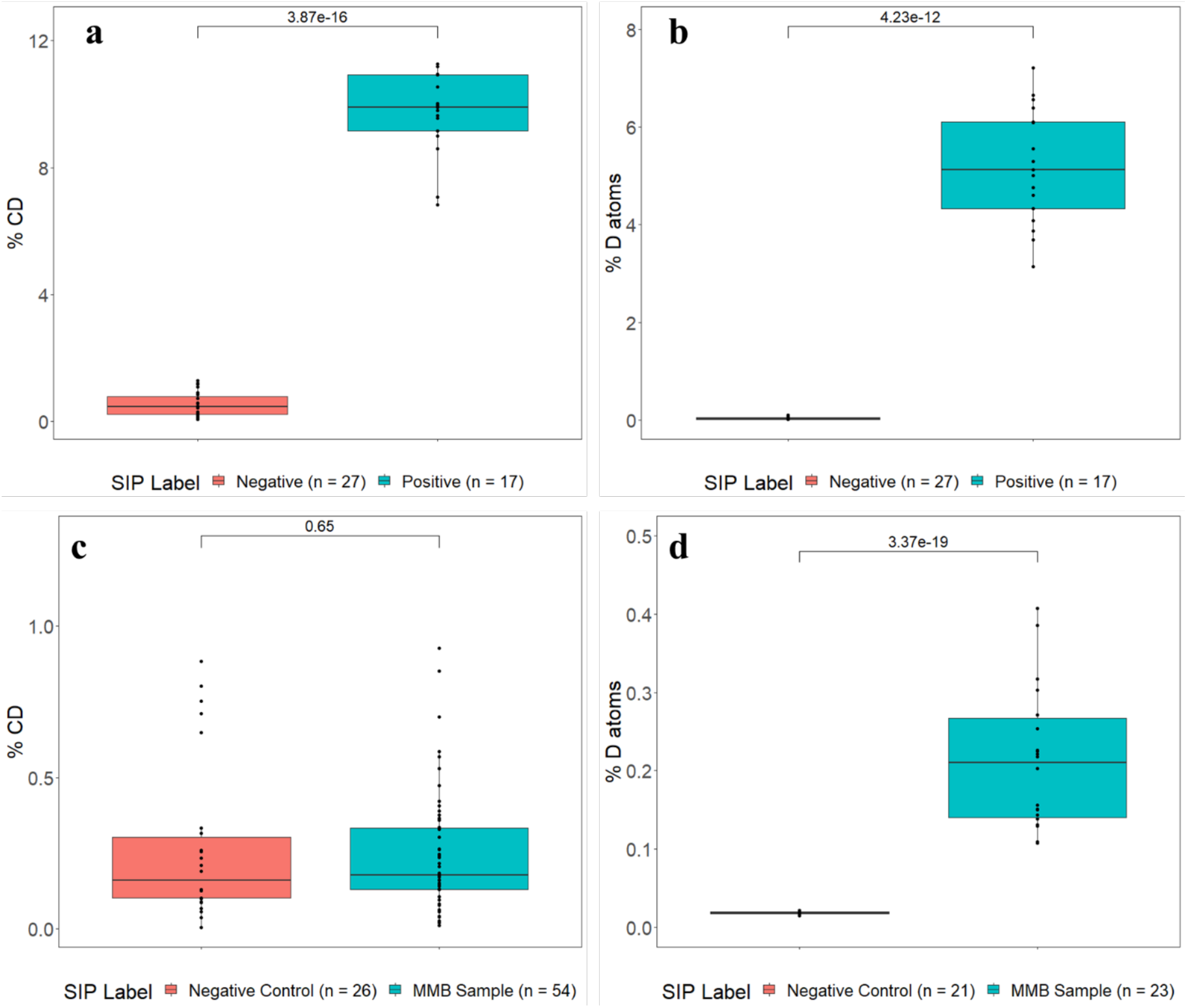
Comparison of D-labeled and non-labeled cells. (a) Analysis of D-incorporation within the mock community calculated from Raman C−H (2,800−3,100 cm^−1^) and C−D (2,040−2,300 cm^−1^) data (p-value = 3.87×10^−16^). (b) Comparison of the NanoSIMS m/z 2/1 (D/H) data for the same cells within the mock community shown in panel (a) (p-value = 4.23×10^−^ _12_). (c) Analysis of D-incorporation within the MMB calculated from Raman C−H (2,800−3,100 cm^−1^) and C−D (2,040−2,300 cm^−1^) data (p-value = 0.65). (d) Corresponding analysis of D incorporated into the cells using NanoSIMS; p-value = 3.37×10^−19^) for the same MMB in panel (c). The black line represents the mean value and individual data points (*i*.*e*., individual cells/MMB) are shown as black dots. Significant differences (p-value shown in plots) were determined by Students t-test.

**Supplemental Figure 4.**
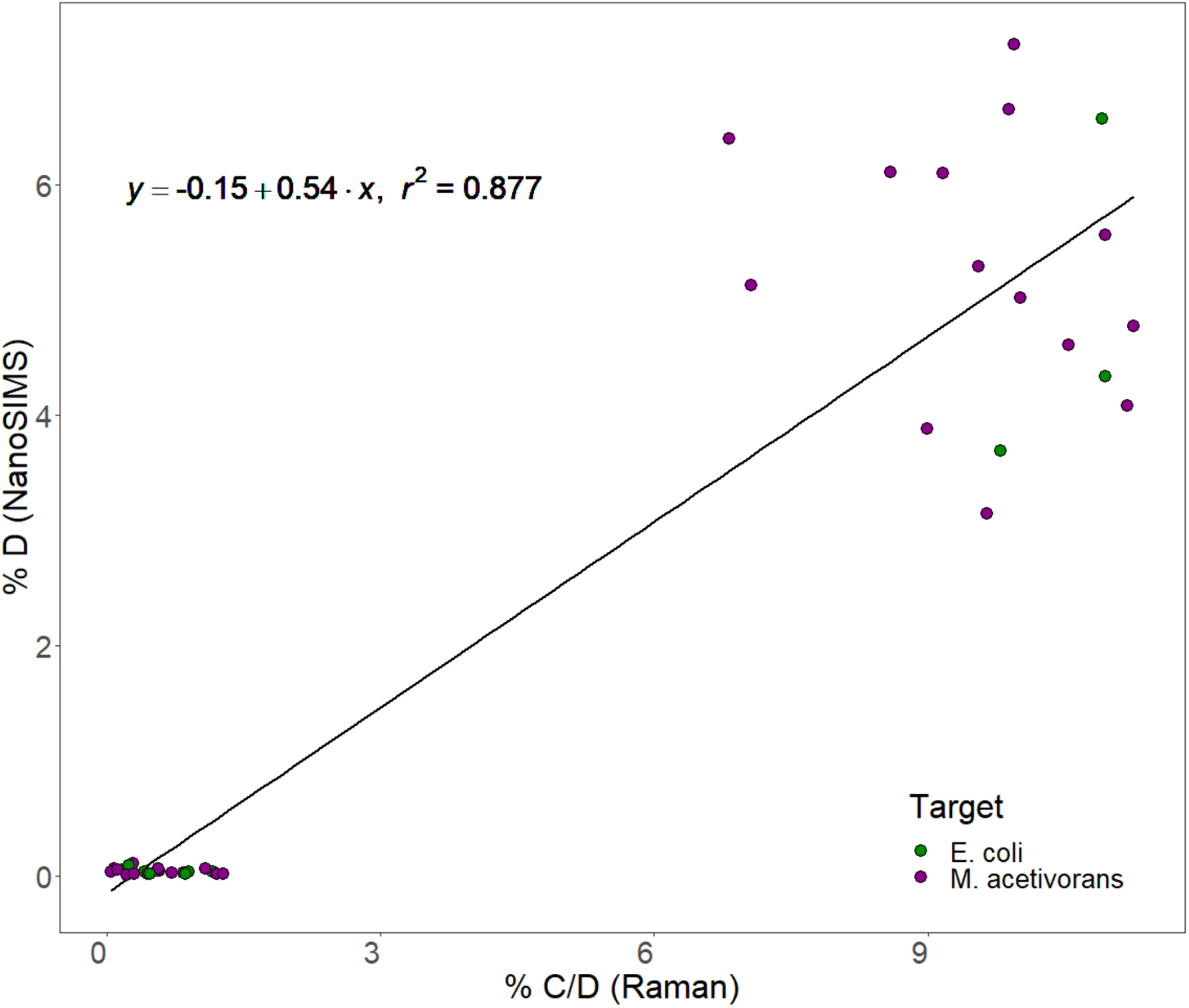
Comparison of Raman and NanoSIMS atom percent of deuterium in the mock community. Both Raman and NanoSIMS were used to detect deuterium within individual cells of the mock community shown in Figure 2. Comparison of data revealed that Raman and NanoSIMS did not yield identical results on individual cells for deuterium incorporation.

